# Postfire recovery of western US conifer forests (1984-2017) using space-borne lidar data

**DOI:** 10.1101/2023.02.18.528502

**Authors:** Nayani Ilangakoon, Jennifer K. Balch, R. Chelsea Nagy, Virginia Iglesias

## Abstract

Coniferous forests account for 78% of the western US forests and store a substantial amount of carbon. Wildfires significantly alter vegetation structure, and hence the forest carbon stock. This study evaluates post-fire vegetation recovery trajectories and rates across the western US using recently launched Global Ecosystem Dynamic Investigations (GEDI) mission lidar data. Three ecoregions studied here, the Pacific Northwest, Southern Rockies, and Northern Rockies, show fire severity and ecoregion specific recovery trajectories for canopy height (CH), plant area index (PAI), and the foliage height diversity (FHD). The recovery trajectories are characterized by an initial decline in vegetation structure (CH, PAI, and FHD) during the first 9-25 years postfire followed by a gain of the structure. Regions of low burn severity can fully recover to the unburned background state within the first three decades while the high burn severity regions may recover in the first century, but only in the absence of fires within this period. The PNW exhibits the slowest recovery rate. According to our results, all three ecoregions feature a loss of growing stock volume (GSV) (−1% - - 48%). Time since fire, fire severity, and altitude were identified as the most significant drivers of postfire vegetation recovery, likely because they integrate the distance to seed source, vegetation composition, and the local climate. Our study suggests that, if fire return intervals become shorter than 50 years, these three ecoregions will have significantly reduced their woody vegetation, hence the carbon stocks. In addition, all the ecoregions studied here exhibit extensive impacts from other disturbances such as beetle invasion. It is therefore important to consider the effects of compound disturbances on vegetation recovery trajectories to infer the future carbon potential in these ecosystems.

## Introduction

Wildfires significantly influence global carbon cycling, ecosystem structure and functioning, and climate change [1]–[3]. Forest fires account for approximately 35% of global fire emissions and can shift these ecosystems, which store a substantial amount of carbon in vegetation and soil [4], [5], from carbon sinks to carbon sources [6]. In recent decades, wildfires across western US forests have increased both in size and frequency [1], [7] due to warming temperatures, human ignitions, and other disturbances [8].

Increased wildfire activity causes substantial forest mortality, carbon emissions, and air quality degradation, and is associated with escalating fire suppression expenditures and property damage [9], [10]. Thus, interest in the impacts of changing fire regimes across the western US and globally has increased, both in terms of short-term effects, such as poor air quality and home destruction, and long-term ecological impact. Most forest ecosystems recover their vegetation/carbon within decades following a disturbance event [11]. However, disturbance legacies can persist longer, causing ecosystem state transitions with altered vegetation structure, composition, and functioning [12]– [14]. Thus, quantifying postfire vegetation state changes and recovery rates is therefore essential to understand how quickly a fire-disturbed ecosystem can initiate vegetation regeneration and recovery to its initial state, the possibility of ecosystem state transitions [14], [15], and how fires influence the global C budget [16], as well as forest structure and functioning.

Most western US forests show mixed-severity (i.e., unburned, low, moderate, and high) fire regimes (Figure 1) with effects ranging from partial damage to complete removal of vegetation within individual fires and across fire events. In addition, over the past two decades, these forests have experienced pronounced drying [17], [18]. High fire severity and increased drought frequency and extent can impede postfire tree regeneration [19], [20]. However, the role of fire severity and climate anomalies in the vegetation recovery trajectory of western US conifer forests is not well understood.

**Figure 1:**
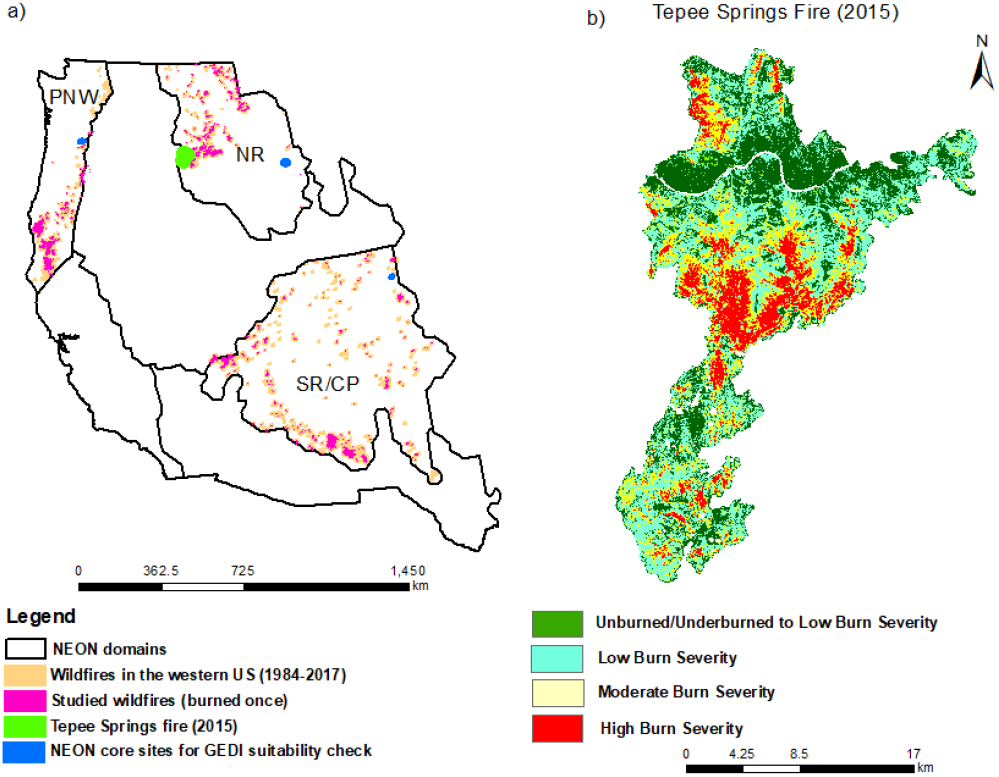
a) Wildfires in the three western US ecoregions as reported in Monitoring Trends in Burn Severity (MTBS; 1984-2017). b) Mosaic of burn severity within a fire (e.g. Tepee Springs fire (2015).

Remotely sensed data have been extensively used to map different fire characteristics including burned area, burn severity as well as postfire recovery [11], [21]. Active remote sensing techniques such as lidar have been used to characterize postfire vegetation structure, aboveground biomass, and carbon stocks at various spatial scales [13], [22]–[24]. In such studies canopy structural metrics such as canopy height, cover, and plant area index (PAI) have been used either with allometric equations to calculate biomass or as a proxy to represent the carbon potential [25].Further, these vegetation structural metrics have been used to infer postfire recovery by measuring the changes in the structure after a fire with respect to the pre-burn state [14]. In this study, we use canopy structural metrics from the first ever high-resolution spaceborne lidar sensor in the international space station, Global Ecosystem Dynamics Investigation (GEDI) [26] along with three fire chronosequences representing three main ecoregions (Pacific Northwest (PNW), Northern Rockies (NR), and Southern Rockies and Colorado Plateau (SR)) to answer the following questions:

1. How quickly does forest structure (i.e., canopy height) recover after fire, and how does the recovery rate vary across ecoregions?

2. To what extent do climatic conditions (e.g. drought occurrence) and fire characteristics (e.g., fire size and fire severity) explain postfire recovery in each ecoregion?

3. To what extent did wildfires in the western US change growing stock volume (GSV) over a 34-year record?

Over the past few decades, all three ecoregions have been heavily influenced by fires of varying intensity and frequency [8]. The fire history, drought, vegetation, and topographic heterogeneity in the study domains provide a unique setting to evaluate postfire vegetation recovery in mountainous, forested ecosystems.

## Methods

### Study area

The study area covers three National Ecological Observatory Network (NEON) ecodomains in the western US: PNW, NR, SR (Figure 1a) [27], [28]. Vegetation cover and composition in the three ecoregions are shaped by climate, elevation, and soil properties. Approximately 56 % of the SR area is mixed forests characterized by ponderosa pine (*Pinus ponderosa*), lodgepole pine (*Pinus contorta*), aspen (*Populus tremuloides*), and pinyon-juniper (*Pinus edulis, Juniperus scopulorum, J. monosperma*, and *J. osteosperma*) [29].

The PNW ecoregion, that has complex land-use and disturbance history is dominated by Douglas-fir (*Pseudotsuga menziesii*) and other true fir species (e.g., *Abies grandis, Abies procera, Abies amabilis, Abies lasiocarpa, Abies concolor*, and *Abies magnifica*). Among many species, ponderosa pine (*Pinus ponderosa*), Douglas-fir (*Pseudotsuga menziesii*), western larch *(Larix occidentalis*), western white pine *(Pinus monticola*), western redcedar (*Thuja plicata*), western hemlock (*Tsuga heterophylla*), grand fir (*Abies grandis*), and lodgepole pine (*Pinus contorta*) are present across the NR ecoregion [29].

### Fire events

We obtained information on 550 fires from the Monitoring Trends in Burn Severity (MTBS) fire occurrence dataset, fire boundary dataset, and burn severity mosaic datasets [30] between 1984 to 2017 (Supplementary Figure 1). We used National Land Cover Datasets (NLCD) between 1985 -2016 to extract fires that occurred only within the Evergreen Forests (NLCD ID = 42) [31], [32]. The NLCD class 42 represents pixels of evergreen forests that are dominated by trees greater than 5 meters tall and 75% of the tree species maintain their leaves all year. Selection of NLCD class 42 data further allows us to use GEDI structure metrics which is only capable of detecting vegetation heights greater than 5 m at higher accuracy [26], [33]. The last land-cover map prior to the fire of interest was used to select the prefire vegetation. From the selected fires, we constrained our study to areas that burned only once. For each fire event, we extracted fire perimeter, fire size, year of the fire, and fire severity conditions (fire severity category, dNBR (Difference Normalized burn ratio) offset, and dNBR thresholds for low, moderate, and high burn severity classes). In this study, we assessed the postfire recovery of fire severity classes 2-4 defined by the MTBS (2: Low Burn Severity, 3: Moderate Burn Severity, 4: High Burn Severity)[30]. Detailed analysis of MTBS data is described in supplementary information.

### GEDI data and the postfire recovery metrics

*W*e used a range of structural metrics that are affected by fire severity, show measurable changes along the postfire period, and are widely used to estimate biomass and carbon. Specifically, we used Global Ecosystems Dynamic Investigation (GEDI) derived vegetation height, plant area index (PAI), and foliage height diversity (FHD) as recovery metrics [33]–[35]. These structural metrics can show measurable changes postfire, and are commonly used to estimate biomass and carbon in lidar based studies. To extract the vegetation metrics, we downloaded all the GEDI_2A v002 and GEDI_2B v002 orbits between May to September of the years 2019, 2020, and 2021e in each ecoregion using NASA Earthdata (https://search.earthdata.nasa.gov/search). As the GEDI waveforms and respective vegetation structural metrics can be affected by snow coverage and by the type of beam (power vs coverage), we used GEDI footprints from power beams between May and September. The data were quality-filtered using a quality flag (= 1) and a sensitivity threshold (> 0.9). After quality filtering, we extracted beam number, shot number, latitude and longitude coordinates, ground elevation, and the RH98 and RH100 metrics from the GEDI_2A product, and cover, PAI, FHD, beam number, shot number, and lat lon coordinates from the GEDI 2B products. The RH (relative height) numbers represent the height that matches with the given percentage of the laser energy from the total laser energy of that particular footprint between the ground and top of the signal. For example, RH100 is the height that matches with the 100% cumulative laser energy of a given GEDI footprint and the height at RH100 is used to represent the canopy top height. Sum of LAI and FHD data from each 5 m interval estimate provided in the GEDI_2B product were used as the PAI and FHD respectively. Beam and shot number information were used to match the variables between GEDI 2A and 2B products. GEDI data preprocessing and extraction were done using modified Jupyter notebooks provided by LPDAAC (https://lpdaac.usgs.gov/news/release-getting-started-gedi-l1b-l2a-and-l2b-data-python-tutorial-series/) and ORNL DAAC (https://github.com/ornldaac/gedi_tutorials) The latitude, longitude spatial coordinates were used to overlay the GEDI footprints over the fire severity and NLCD maps to extract fire severity data for each footprint based on their geolocation. We developed postfire recovery trajectories based on canopy height, PAI, and FHD only. We further tested the suitability of GEDI structural metrics to assess the postfire vegetation recovery in regional scale and details of this can be found in the supplementary information and Supplementary Figure 3.

### Climate variables

The following variables were used in the analyses: mean annual precipitation (mm), standardized precipitation evapotranspiration index (SPEI06 - SPEI calculated using 6 months differences between precipitation and potential evapotranspiration), relative humidity, and vapor pressure deficit (VPD; kPa). Climate data layers representing each variable mentioned above between 1984-2017 were extracted from the Climatic Research Unit database, which provides the data at 4-km spatial resolution [36], [37]. To facilitate across-fire climate comparisons and account for their impact on postfire recovery, the selected climate variables were converted to postfire climate anomalies. We first log transformed the precipitation data to correct the skewness of the original precipitation data. Then we calculated the z-score or the climate anomaly for each climate variable (precipitation, SPEI and VPD) following below equation [38] :

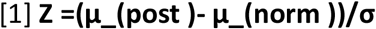

where Z, μ_(post), μ_(norm), and σ are the Z statistic, postfire mean climate, mean climate normal between 1984-2017, and standard deviation of the climate normal between 1984-2017, respectively. To calculate the mean postfire climate for each fire, we extracted values from each climate variable from one year after the ignition year till 2017. If more than one climate pixel is found within a single fire, we use the average, so that only one value per climate variable per fire is produced.

### Building postfire recovery trajectories

We built a postfire recovery trajectory for three chronosequences, PNW, NR, and SR/CP (SR hereafter). Our study focused on fires where time since fire is between 1-33 years (1984-2017) to evaluate the recovery rates and trajectory. MTBS pixels were used as the sample unit in this study. Explanatory variables were the GEDI footprint based (25 m) structural metrics (described above). We selected GEDI footprints surrounded by a single severity class up to 45 m from its defined coordinates, to avoid footprints that fall into different severity classes due to geolocation errors associated with GEDI data (Approximately 10 m). We randomly selected a maximum of 200 GEDI footprints from the surrounding area of each fire with no known burn history to represent the unburned (control) state. We used all the GEDI footprints within the class of “42” of the national land cover datasets (NLCD) to select the same land-cover type for the unburned areas. We excluded the area within 100 m from the fire perimeter to minimize errors associated with detecting the fire boundary. Percent change of CH, FHD, and PAI values of the burned regions with respect to unburned background were used as the recovery metric.

Percent change of CH, PAI or FHD values of the burned regions with respect to unburned background were used as the recovery metric and were evaluated as a function of fire (severity, time since fire), site (elevation, distance to unburned area), and climate (precipitation, temperature, drought as represented by SPEI, and relative humidity). We categorized time since fire (and thus vegetation age) in 3-year, 5-year,7-year, and 10-year intervals from 1984 till 2017. The 3-, 5-, 7-or 10-year intervals were defined to capture distinct growth episodes along the recovery trajectory (e.g., initial loss, establishment, rapid recovery and maturity phases as well as transitions between those phases). We used a one-way ANOVA to identify the interval that minimized within-group differences while maximizing among-age group variations in the chronosequence. Detailed information of grouping fires into different time since fire intervals and the results of One-Way ANOVA can be found in the supplementary information and Supplementary Figure 4.

To identify the shape of the recovery trajectory (Q1), we fitted nonlinear models of Gompertz, Chapman‐Richards, Michaelis‐Menten, and polynomial functions to the original GEDI metric points using R package “growthmodels”[4]. Our selection of non-linear models was based on visual inspection of the data distributions supported by evidence of non-linear chronosequences of carbon recovery in a range of ecosystems [39], [40]. Model performance was evaluated with Akaike Information Criterion (AIC) [41]and standard residual errors. In each ecoregion, the model that minimized both estimates was used to inform on recovery rates and time required to recover to the background state.

To explore the role of fire size, climate, and fire severity on postfire vegetation recovery, we used Generalized Additive Models (GAMs). Given the non-normality of the data and the potential for linear and nonlinear effects of the predictors on recovery trajectories, data-driven models that allow the specification of the distribution of the response variable and the link function such as GAMs [42] are optimal tools for association estimation. To assess the relative influence of the explanatory variables on GEDI-derived structure metrics (recovery trajectories), we followed a forward stepwise selection process. That is, starting from a null model, we added predictors one at the time and compared the performance of the respective models based on their AIC. Structural metrics from undisturbed sites were used as the reference state to estimate the change in vegetation structure at each time step. The data was processed in R (R studio) using packages “nlme” and “drc”, and “gam”.

### Growing stock volume (GSV)

As we do not have wood density estimates across the three ecoregions, we did not estimate the biomass or carbon. However, to infer the carbon gain postfire, we modeled the growing stock volume using GEDI derived canopy heights and the plant area indices as GSV show a linear relationship with tree biomass and carbon [43], [44]. GSV in this study is calculated as the product of canopy height and the PAI after 33 years of fire. The PAI represents the projected ground area from all canopy materials within the GEDI footprint[45]. In this study, we first counted the number of pixels in each fire severity class within each ecoregion using ArcMap 10.8. Then, we multiplied the modeled canopy height by modeled PAI after 33 years of the fire at each severity class. This is the mean per pixel canopy volume we expect from each severity class after 33 years of the fire. Then, we multiplied these estimated canopy volumes by the number of pixels of each severity class at each ecoregion to get the total growing stock volume after 33 years of a fire. We further calculated the uncertainty of canopy volume estimates using 95 % confidence interval estimates of canopy height and PAIs from our recovery trajectory models. In addition, using average canopy height and PAIs together with their uncertainties from the unburned regions was used to calculate the growing stock volume if this total burned area of each ecoregion was not burned.

## Results

### Canopy height recovery across ecoregions

The mean canopy heights of the unburned regions are 20.71 m, 27.71 m, and 10.36 m for NR, PNW, and SR respectively. One-way ANOVA confirmed that mean CH among 10 year interval groups (based on time since fire from 1984 till 2017) are significantly different than within group variations and thus were used to build the recovery trajectory (Supplementary Figure 3). Both SR and PNW do not show a clear change in canopy heights at 3-, 5-or 7-year intervals while the NR started showing a clear declination of CH followed by a gain with time since fire even at 5 year intervals. A second order polynomial function provided the best fit model for fire recovery trajectory with the original data (Grouped by 10 year intervals) for all three ecoregions (Figure 2 and 4). Canopy height shows an initial decrease followed by an increase with time since fire. The magnitude of the increase or decrease varied with respect to fire severity and ecoregion (Figure 2). In the PNW, overall change in postfire canopy height closely follows the canopy heights of the unburned curve (Figure 2a). However, when considering the fire severity, a larger decline in percent canopy height is observed with both the moderate (−10%) and high severity (−25%) regions (Figure 5a). These decreased percent canopy heights remain almost consistent with very little change over the next two decades and start to decline further around 25-30 years postfire. The percent canopy heights of the low burn severity regions slowly increase over the first 33 years postfire. There is no canopy height gain observed in both moderate and high burn severity regions in the PNW over the 33 years studied. A greatest decline in overall canopy height postfire is observed in the NR (Figure 4b). Detailed analysis shows a decline in canopy height in all three severity classes in the first 20-25 years in the NR region. The largest decline in canopy height is observed in the high burn severity regions (−52%) (Figure 2b). The low burn severity regions also show a considerable declination of canopy height (−23%) after 22 years of a fire. However, only the low burn severity regions in the NR reach 100% recovery after around 35 years of the fire. Similar to the PNW, the SR also shows a little to no change in overall canopy height postfire over the 33 years of records (Figure 2c). However, the analysis based on fire severity shows a mild decline in percent canopy heights (−8%) followed by a very slow gain in high burn severity regions after 18-20 years in the SR. Both the low and moderate burn severity regions show less than 5% canopy height decline (around 18 years postfire) and fully recover within the first 3 decades (Figure 2c).

**Figure 2:**
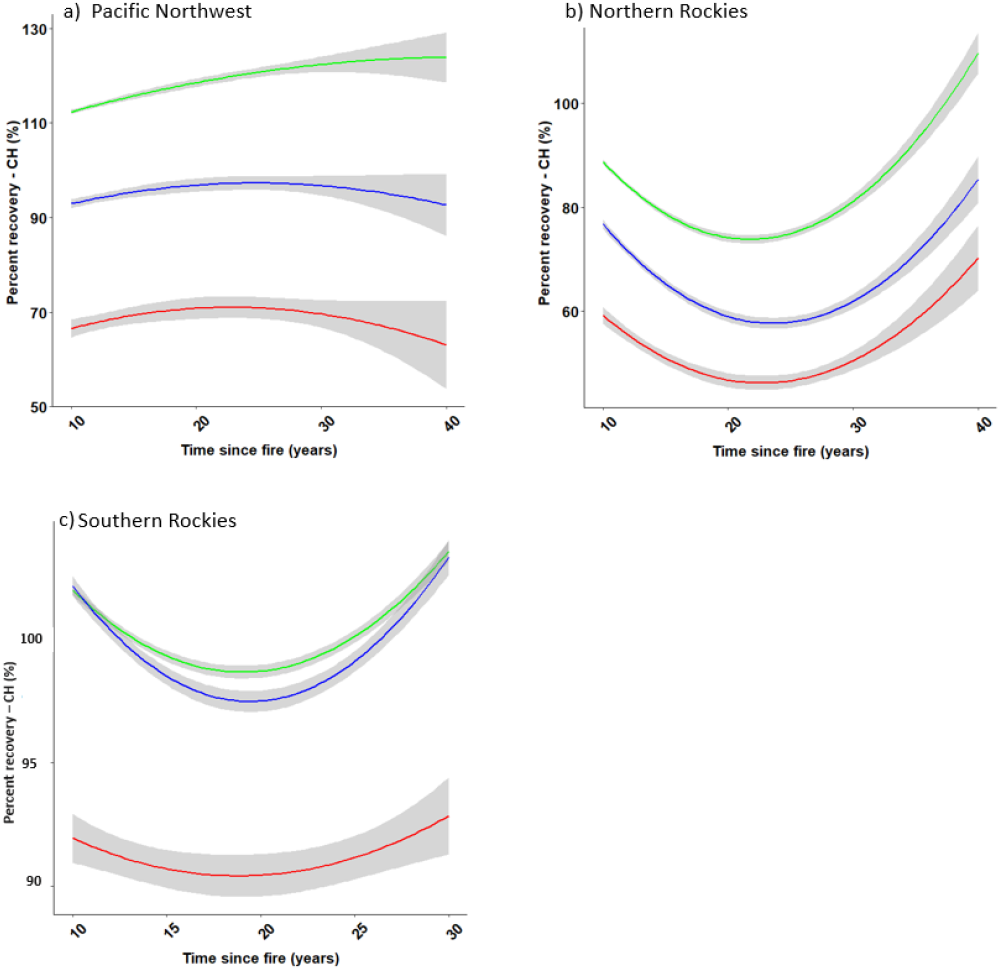
Post-fire percent canopy height (% CH) recovery trajectories with burn severity in a) PNW, b) NR, and c) SR. Green: Low severity, blue: Moderate severity, red: high severity. The shaded areas around solid lines represent the 95% confidence interval.

Overall, the largest decline in canopy height is depicted in areas with high burn severity in all three ecoregions (−8 % - 52% of the unburned state), while the lowest decline was observed in the low burn severity areas (0% - 22.2% of the unburned state).

All three ecoregions show percent foliage height diversity change with time since fire similar to that of their percent canopy height change (Figure 3). PAI slowly recovers in all three ecoregions. The PAI of low burn severity regions in all three ecoregions fully recovers within the first three decades (Figure 4). The PAI of both the moderate and high burns severity regions of the PNW and the NR show continuous recovery. However, the PAI of high burn severity regions in the SR come to a steady state after 22 years of the fire (Figure 4c). Overall, the PNW shows the slowest recovery rate while the NR the fastest recovery rates. Though we did not extrapolate the trajectories, the trends in high severity trajectories in all three ecoregions suggests that they may not recover to the background state even within the next two decades (∼ 50 years postfire).

**Figure 3:**
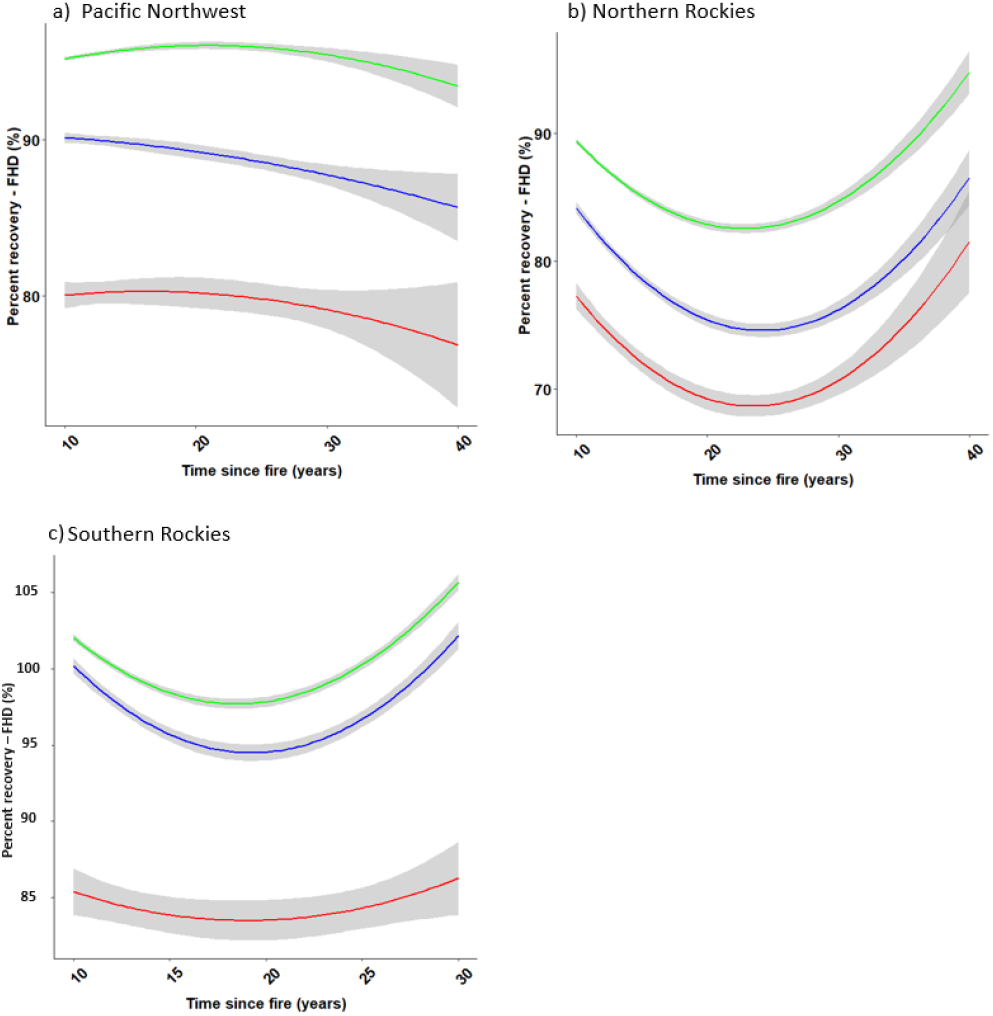
Post-fire percent foliage height diversity (% FHD) recovery trajectories with burn severity in a) PNW, b) NR, and c) SR. Green: Low severity, blue: Moderate severity, red: high severity. The shaded areas around solid lines represent the 95% confidence interval.

**Figure 4:**
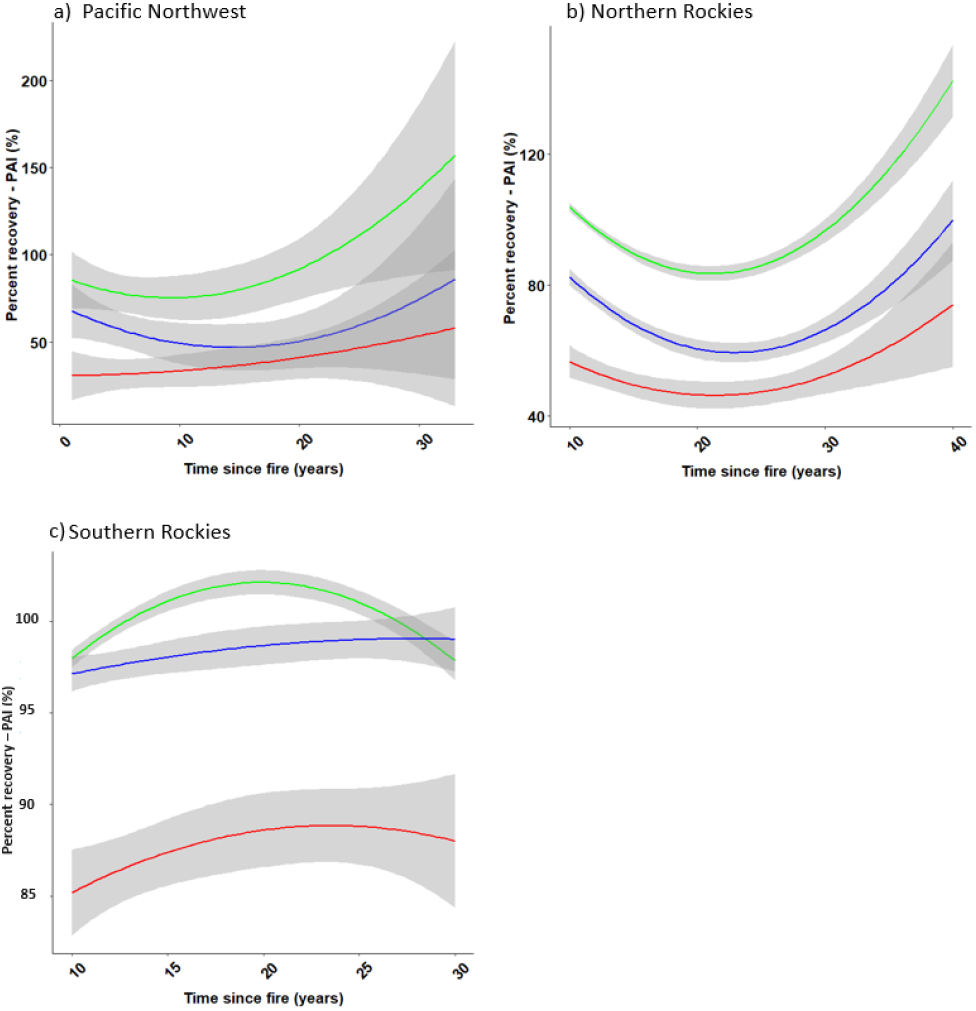
Post-fire percent plant area index (% PAI) recovery trajectories with burn severity in a) PNW, b) NR, and c) SR. Green: Low severity, blue: Moderate severity, red: high severity. The shaded areas around solid lines represent the 95% confidence interval.

### Effect of climate and site characteristics on fire recovery trajectory

When considering all three ecoregions together, time since fire alone explained 50% - 60% of the CH recovery. While the time since fire shows a non-linear effect on postfire recovery, fire severity, elevation, distance to undisturbed areas and fire size show a linear effect with the postfire canopy height gain (Figure 5). When considering each ecoregion as a separate unit, the important controlling factors were significantly different. In the PNW region, other than time since fire, elevation, fire severity, distance to undisturbed state significantly controlled the postfire canopy height gain. Based on our results, postfire recovery in PNW decreased when the elevation, fire severity, and the distance to undisturbed areas increased.

**Figure 5:**
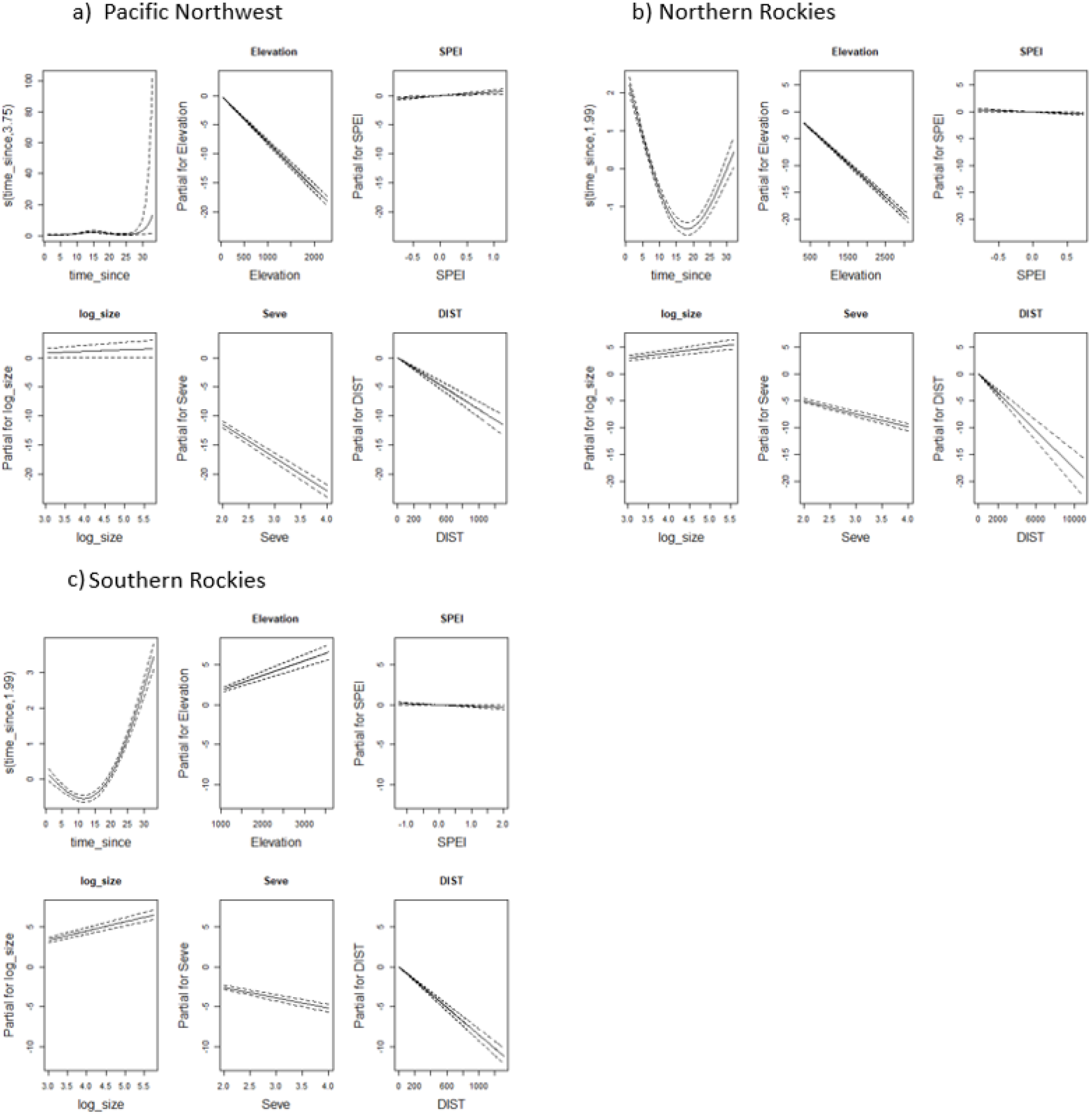
Controls of postfire canopy height recovery in the three ecoregions. a) Pacific Northwest, b) Northern Rockies, c) Southern Rockies. The x-axs are variables. The y-axs represents the partial effect of each variable. The area between dash lines indicate the 95% confidence intervals. ariables : Time since: years since the fire to 2018, Elevation: ground elevation above mean sea level, Seve: burn severity, DIST: closest distance to unburned region, log_size: burned area converted to log scale, SPEI: Standard precipitation evapotranspiration index

Similarly, the recovery trajectory of the NR region is also controlled by time since fire, fire severity, and elevation. Fire size showed a mild positive effect on postfire recovery in the NR region, with larger fires associated with faster gains of canopy height. Same factors control the postfire recovery trajectory of the SR region as of the NR. None of the ecoregions show a canopy height recovery trend with respect to the drought occurrence (aka SPEI).

### Postfire growing stock volume (GSV) in the western US

Estimated GSVs show a significant reduction compared to the unburned background state in all three ecoregions (Figure 6). The lowest percent GSV is observed in the NR (1.1%) while the highest GSV is reported in the low severity burn areas of the SR (48%). Overall, the SR region shows a higher GSV (23 - 48%%) compared to the other ecoregions. However, none of the ecosystems recover to its unburned vegetation volume (and potentially the biomass or carbon) after 33 years of the fire, even in the low burn severity regions.

**Figure 6:**
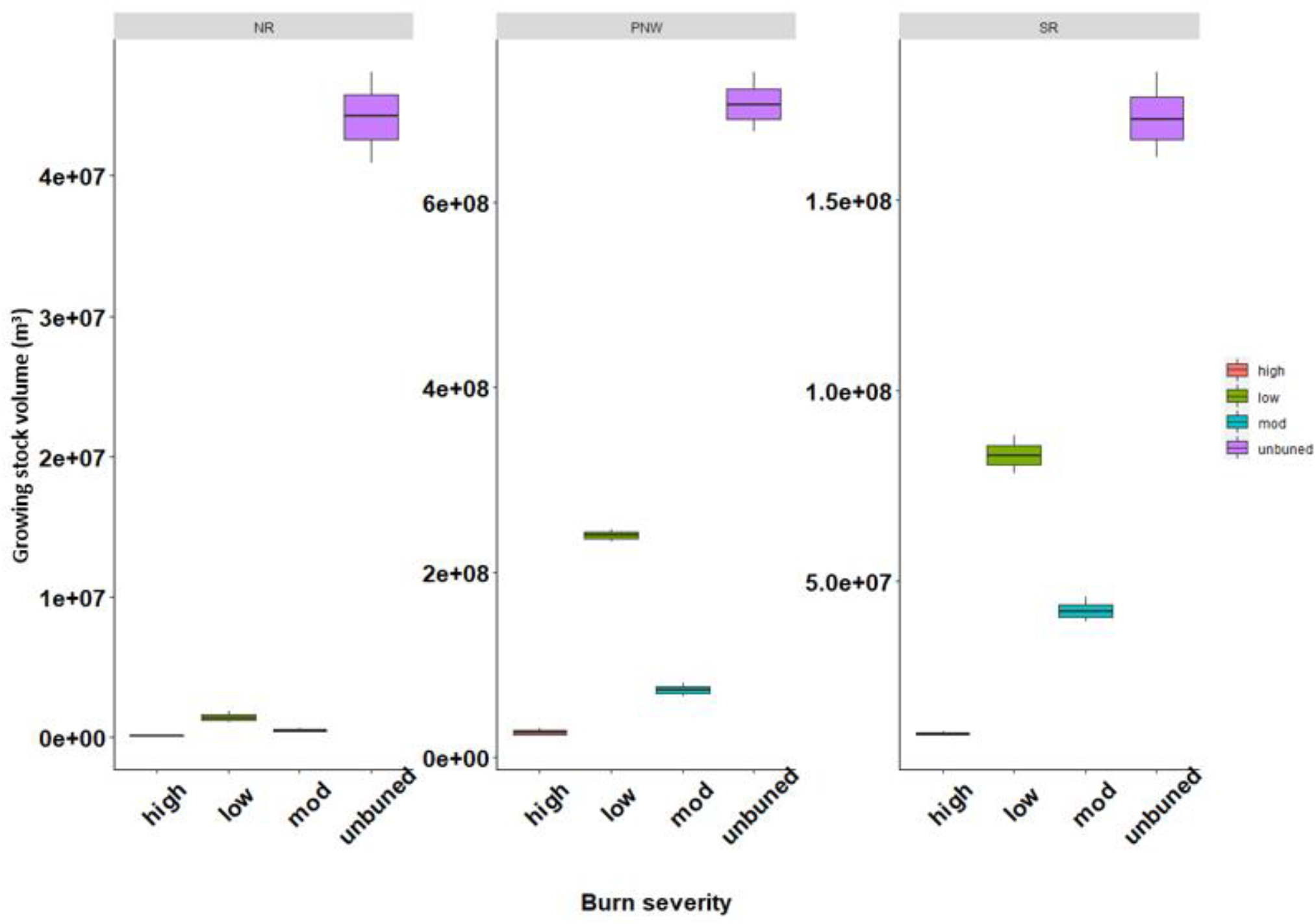
Growing stock volume after 33 years postfire calculated using modeled vegetation recovery in each ecoregion in the western US. Please note the scale difference in the y axis (scale differences were used for visualization purpose)

## 4. Discussion

### Postfire vegetation recovery trajectory

Fire driven tree mortality followed by vegetation regeneration and succession contribute to the future carbon potential of an ecosystem. Therefore, quantification of vegetation regeneration, succession timing, and trajectory after a fire is critical to estimate the ecosystem carbon stock and the global carbon balance, especially under changing climate. We developed postfire vegetation recovery trajectories within three main western ecoregions. We used the newest spaceborne lidar (i.e., GEDI) based vegetation structural metrics over 550 fires across the western US between 1984-2017. We used the newest spaceborne lidar (i.e., GEDI) based vegetation structural metrics over 550 fires across the western US between 1984-2017. This study evaluates canopy height, plant area index, foliage height diversity, and growing stock volume changes within the first three decades of the postfire period as an initial estimate of the vegetation and carbon reserves of three main western US ecoregions. Further, this study evaluates the key drivers of the vegetation recovery trajectory and the carbon potential of western US ecoregions after a fire.

We use canopy height as the primary indicator of forest recovery as it is widely used to estimate biomass and carbon stock as well as tree mortality from disturbances [23], [34], [38]. According to our results, the near term (three decades postfire) canopy height recovery is represented by a nonlinear, asymmetrical “U” shape curve in all three ecoregions, likely representing the initial and delayed tree mortality followed by vegetation regeneration and succession [10], [46]. While some postfire regeneration studies use vegetation density as a regeneration metric [45], [47], we used vegetation growth (gain of canopy height and growing stock volume) as a proxy for carbon gain postfire as our main focus to assess carbon recovery potential in these three ecoregions. The rate of change of CH in moderate and high severity burned areas is an indicator of snag fall as well as the percentage of snags to the surviving trees. In addition, we used Plant Area Index and foliage height diversity to infer the vegetation cover and density gain postfire.

Each time since fire intervals (10-year intervals or age groups) in our study represents fires that are spatially distributed within each ecoregion, however, we did not have an equal number of fires for each of these 10-year intervals. Though we selected burned areas within evergreen forests as the prefire dominant vegetation community, each burned and unburned area may contain other vegetation including deciduous trees (e.g., aspen) of different growth rates. The shape of the postfire recovery trajectory largely depends on what species survived and regenerated under the disturbance. The variation of canopy heights within age groups in each ecoregion can be attributed to this heterogeneous vegetation composition and structure. Though the percent canopy height change remains steady throughout the first three decades in the PNW, the increase in both the percent FHD and PAI may represent the vegetation recovery likely due to the growth of understory. The slow decline in mean canopy height in all severity classes in the PNW around 25 years postfire may be due to other compounding disturbances which can cause tree mortality (e.g. beetle invasion) or could be due to small fires that were not included in the MTBS database. As we had only a few fires in the 1980s, even if a single fire is affected by another disturbance, a larger difference in the trajectory can be caused. In the PNW, Whitlock et al., 2015 [48] showed that the rate of forest regeneration can take multiple pathways for initial cohort development, and all these pathways reach canopy closure in about 4 decades. The extrapolation of the recovery trajectory in our study, however, does not show a full canopy height recovery in the PNW even within the first 5 decades. Further, our age groups (3-, 5-, 7-, or 10-) and the time period (33 years) may not be representative to capture the tree mortality and growth episodes in the PNW.

In the NR ecoregion, our results are consistent with those of Stevens-Rumann et al., 2020 [49], as they show reduction of standing trees during the first 24 years postfire from both stands replacing (high severity) and non-replacing fires (low to moderate severity). The greater the distance to seed sources, can further reduce the postfire vegetation establishment [50]. The higher recovery rates associated with larger fire sizes may represent faster vegetation regeneration due to increased light resources and nutrient availability. In this study, however, we did not consider fire refugia size and spatial distribution, which may have an influence here promoting faster vegetation growth regardless of the fire size. Consistent with our findings, previous studies show that the fire severity within each stand can drive post-fire vegetation recovery patterns in evergreen forests with low burn severity regions showing a faster recovery while the high burn severity regions show the slowest recovery [51], [52]. In addition, stand-specific research in conifer forests yielded similar results [20], [49], confirming the influence of site characteristics such as topography (elevation) on postfire recovery. The NR region is mainly characterized by Ponderosa pine (*Pinus ponderosa*)[49]. As a fire-resistant species, ponderosa pine tends to dominate in the NR region over all other species. In addition to the above species, Quaking aspen (*Populus tremuloides*), a fire-tolerant deciduous species, can dominate burn scars [53], [54] and replaces conifer trees that were killed by beetle invasion. However, in high severity fires, there will be limited seeds for regeneration leading the ecosystem to an open environment with different vegetation abundance and spatial structure. The larger decline of canopy height in high severity fires of the Northern Rockies shown in our study could be due to these lack of seed sources for both deciduous and conifer tree species. This large decline in canopy height could eventually reduce the carbon storage and carbon potential in these ecosystems. Research further show that fire suppression during the past century in the NR has increased the fuel load in low elevations while in high elevations, reduced snow packs have increased fire vulnerability [49]. This, coupled with the large influence of topography in postfire recovery, emphasizes the importance of future studies that evaluate how topography-related features such as slope, aspect, and elevation, shape postfire vegetation recovery in the NR region.

The SR ecoregion comprises ponderosa pine, piñon-juniper woodland, high-elevation subalpine forests as well as aspen, hence creating a variable fuel complex. The fast rate of regrowth in the low and moderate burn severity regions in the SR could be due to an accelerated growth of existing vegetation due to the clearing (reduced competition to light). Our results show that in this ecoregion, fire severity and elevation show a larger contribution to the vegetation recovery trajectory shape than in the other two ecoregions. In agreement with our results, [20], [55] also showed that postfire regeneration can fail tremendously with larger fires and droughts, especially in ponderosa pine dominated regions.

In this study, we considered areas that burned once, and did not consider compound disturbances. However, the frequency and magnitude of fire disturbance together with other disturbances defines the distribution of vegetation composition in these ecoregions [55]. Evidence shows that tree mortality from increased mountain pine beetle invasion further reduces the carbon stock of fire-disturbed ecosystems in the PNW region. Beetle impacted tree mortality also drives changes in vegetation structure and composition and the carbon stock in the NR [56]–[58]. Increased fire frequency, larger fires, and pine beetle invasion together can induce ecosystem state changes, converting forests into open vegetation and thus increasing the carbon vulnerability in the western US. Hence, future studies that consider the impact of compound disturbance in post-disturbance vegetation recovery across ecoregions are essential to estimate the true carbon potential of these ecoregions.

Fire is a powerful ecological and evolutionary force that regulates carbon and nutrient cycling and informs about the global carbon vulnerability. Vulnerability in this study is considered with respect to each ecoregion and its potential to recover after a single fire. Our results suggest that none of the ecoregion will recover to full canopy volume within the first 50 years postfire. Only the low burn severity regions will gain canopy heights similar to the unburned state within the first 5 decades, however, due to the differences in plant area index, canopy volume still be well below the unburned state indicating these burned systems as long-term carbon sources. If the fire return intervals are shorter than this 50-year period, these ecoregions will potentially convert to alternative states. According to fire histories and model analyses provided in the LANDFIRE products [59], the Northern Rockies has a longer fire return interval (∼ 137 years). The fire return intervals in the PNW is approximately 25 years. Hence, PNW regions are at higher risk of losing carbon potential than the other two ecoregions. The SR shows a fire return interval of around 46 years. These fire return intervals are the median return intervals from the LANDFIRE FRI of all fires from Conifer and hardwood-Conifer dominant regions of these three main ecoregions. However, all three ecoregions are significantly impacted by other disturbances that cause tree mortality and increased fire frequency. The occurrence of other disturbances in these fire-disturbed areas further reduce the ability of these forests to completely re‐accumulate the biomass to prefire levels. Hence, it is important to evaluate the occurrence as well as how these other disturbances influence postfire vegetation recovery.

### GEDI for forest recovery estimates

The GEDI provides high-quality measurements of forest vertical structure in temperate and tropical forests between latitudes 51.6° N & S at a footprint resolution of 25 m. As GEDI footprint-based biomass estimates have not completely been released, we constrained our study to structural metrics. The GEDI footprint-based canopy height, canopy foliar profiles and Plant Area Index (PAI) are critical parameters to understand canopy vertical variations within and across ecosystems based on structure, composition and function. As we used GEDI-estimated CH as the primary recovery indicator, GEDI height retrieval accuracy can also impact our results. The larger pulse length (15.6 ns) of GEDI is only capable of detecting vegetation heights greater than 5 m without correction for the influence of the pulse length or ground slope. With the fastest recovery rate (0.74 m y^-1^ in the SR ecoregion), it takes more than 7 years for the young vegetation to be captured by GEDI. Hence, we assume that the initial decline in canopy height in our trajectories is represented by the snag fall while the start of canopy height gain is represented by any vegetation greater than 5 m. We emphasize that future studies integrating high-resolution airborne lidar or UAS based point clouds to better understand the initiation of postfire vegetation regeneration and succession is critically important.

## Conclusions

Forests are important drivers in global carbon dynamics, storing a substantial amount of carbon in vegetation and soil. Wildfires across western US conifer and mixed conifer forests are increasing in size, frequency, and number due to warming temperatures, human ignitions, and other disturbances. Estimates of postfire carbon change and recovery fulfills a dire need to understand how increasing fire activity can influence regional and global carbon budgets. Using GEDI estimated vegetation structure, our study shows that post-fire vegetation recovery rates in the western US are ecoregion specific, with the fastest recovery in the Southern Rockies and the slowest recovery in the Pacific Northwest forests. The recovery rates are controlled by time since fire, fire severity, and elevation gradient with different magnitudes depending on the ecoregion. In addition, the recovery of vegetation in these forests are sensitive to the distance to undisturbed regions and fire size emphasizing the accessibility to seed sources. Our results further emphasize that none of these ecoregions recover their full canopy volume within the first 5 decades postfire indicating long term reduced carbon stocks.

## Data availability statement

The data that support the findings of this study are available upon reasonable request from the authors.

## Authorship roles

N.I and J.K.B conceptualized the study. N.I., J.K.B., and V.I., processed and analyzed data. N.I, J.K.B., R.C.N., and V.I. wrote the paper.

