## Supplementary figures for "Postfire recovery of western US conifer forests (1984-2017) using space-borne lidar data"

### Supplementary Materials

#### MTBS data pre-processing

a)

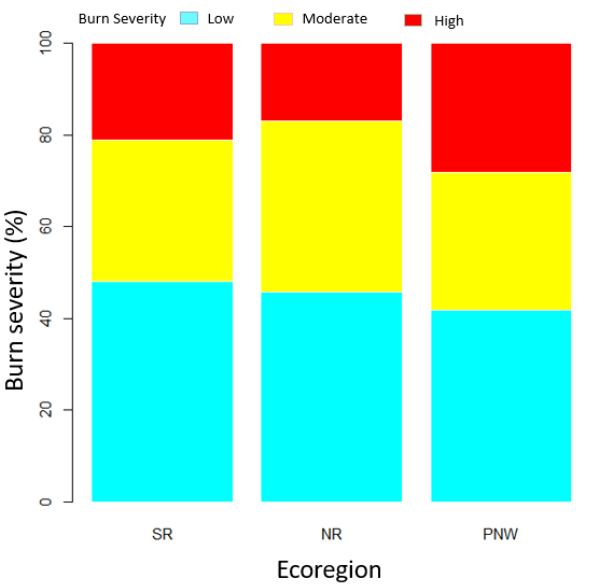


Supplementary figure 1a: Percentage of pixels with each burn severity category in PNW, SR, and NR ecoregions.

We downloaded fire occurrence dataset, fire boundary dataset, and burn severity mosaic data layers between 1984 to 2017 from the Monitoring Trends in Burn Severity (MTBS) website (https://www.mtbs.gov/). We used fire year, fire type (we selected only “wildfires'') and burned area polygon data in ArcMap 10.8 (Esri, West Redlands, CA, USA) to extract wildfires that burned only once between 1984-2017 in each ecoregion (PNW, NR, and SR). Then, overlaying GEDI footprints data over selected fire data, we extracted fire information (fire size, year of the fire, and fire severity conditions defined by the MTBS (2: Low Burn Severity, 3: Moderate Burn Severity, 4: High Burn Severity) Supplementary figure 1 .

b)

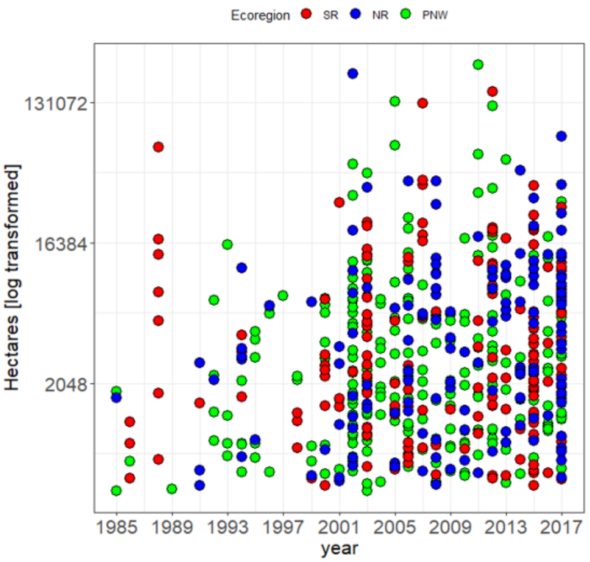


Supplementary figure 1b: Fire size and ignition year distribution of fires used in this study. There are more fires in the later decades than the 1984- 2000 period in each ecoregion.

We further extracted the NLCD class IDs for each GEDI footprint from all the land cover data sets available between 1985 - 2016. Using these MTBS, NLCD, and GEDI data, we extracted all the GEDI footprints that showed only one fire (aka Burned once) and are within NLCD class ID 42. The last land-cover map prior to the fire of interest was used to select the prefire vegetation class. MTBS fire severity data are developed on a fire basis and are not directly comparable across fire events [54]. Hence, we subtracted dNBR offset values from each severity class dNBR threshold to create offset corrected dNBR threshold for each severity class to analyze the distribution of modified dNBR thresholds across fire severity class and ecoregions (Supplementary figure 2). We observed that after offset correction, fire severity classes are well separable in each ecoregion and used this information in postfire vegetation recovery estimates and result interpretation based on fire severity.

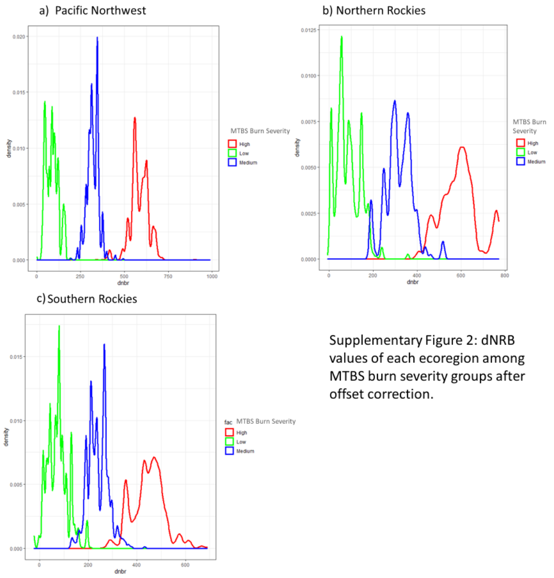


Supplementary Figure 2: dNRB values of each ecoregion among MTBS burn severity groups after offset correction.

### Suitability of GEDI data for postfire recovery estimates

We analyzed the suitability of GEDI canopy height to estimate vegetation heights in the studied ecoregion. To do this, we used National ecological Observatory Network (NEON) based 1 m canopy height products developed using airborne lidar data (ALS hereafter). To represent each ecoregion, we downloaded NEON Canopy height data developed in the 2019-2021 period from three core sites (Yellowstone Northern Range (YELL), Niwot Ridge Mountain Research Station (NIWO), and Wind River Experimental forest (WREF)) from each NEON domain we studied. The YELL , NIWO, and WREF represent Northern Rockies, Southern Rockies, and Pacific Northwest ecoregions respectively. We aggregated NEON 1m canopy height data into 25 m spatial resolution using max, mean and median. We also created a circular buffer of 25 m diameter around each GEDI footprint using center lat and lon coordinates. Then we extracted ALS vegetation heights for GEDI footprints that cover more than 80% of the ALS 25 m pixel. We used this 80% overlap between buffered GEDI footprint and ALS 25 m pixel to minimize the impact of geolocation errors associated with both GEDI and ALS data. While all three ALS based height estimates show correlation above 50%, ALS based median heights showed the highest correlation with the GEDI estimated heights (r = 53%) (Supplementary figure 3a).

As GEDI biomass data release is not completed and has less biomass data over our study region, we used GEDI based canopy heights in this study. Though GEDI based biomass data are developed using GEDI RH metrics, we first wanted to test the suitability of canopy height usage for our study. We downloaded available biomass data over our study region and processed the data using a modified Jupyter notebook based on ORNL DAAC (https://github.com/ornldaac/gedi_tutorials/blob/main/2_gedi_l4a_subsets.ipynb). We used aboveground biomass density data at default setting with *l2_quality flag> 0.9, and l4_quality_flag> 0.95* in this study. The linear regression analysis showed that the max canopy height can predict GEDi biomass with R2 = 0.9 (Supplementary figure 3b). f

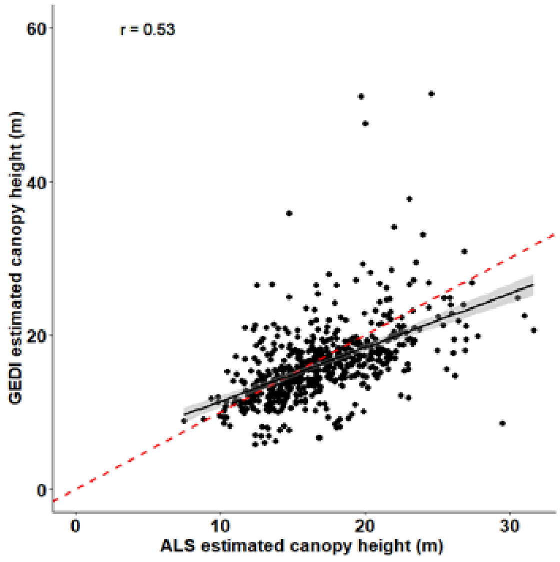


Supplementary figure 3a: Correlation between airborne lidar and GEDI lidar derived canopy heights in the western US evergreen forests.

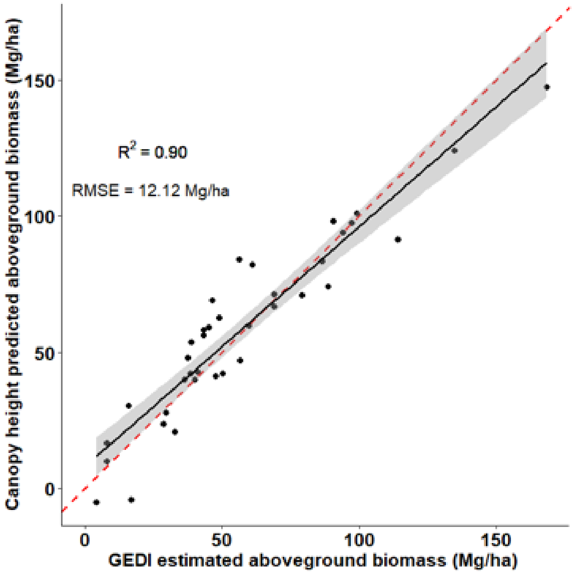


Supplementary figure 3b: Linear regression between GEDI estimated biomass and GEDI canopy heights derived biomass in this study

**Grouping fire data to represent tree mortality and regeneration episodes**

To select the best group of postfire years to clearly represent the initial loss, establishment, rapid recovery, and maturity phases as well as transitions between those phases we grouped postfire canopy heights into 3-, 5-, 7- and 10-year intervals. Supplementary figure 3 shows the One-Way ANOVA results computed as Tukey Honest Significant Differences using R. According to the figure below (Supplementary figure 4), intra group means are lower than the neighboring group means. The histograms of canopy heights at each group is shown in Figure 4.

Pacific Northwest

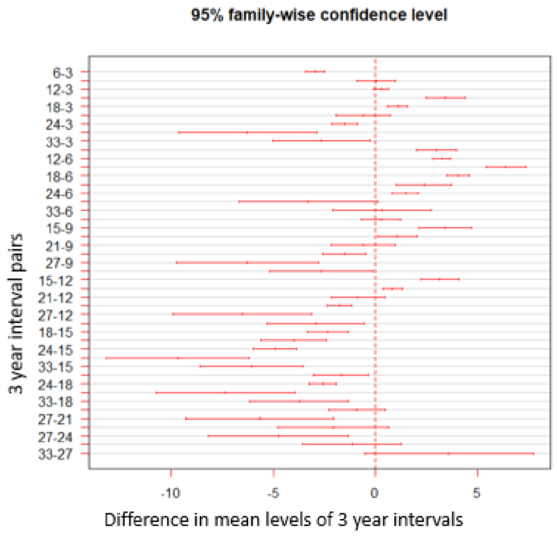


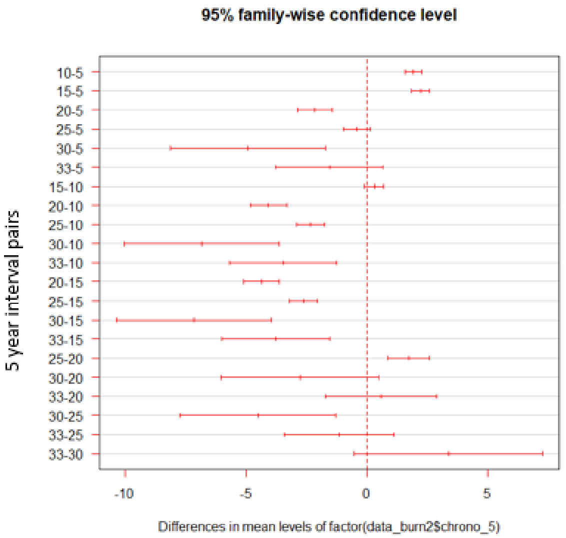


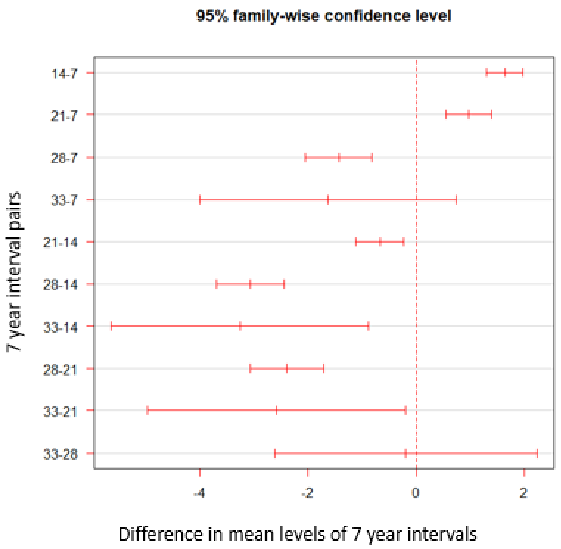


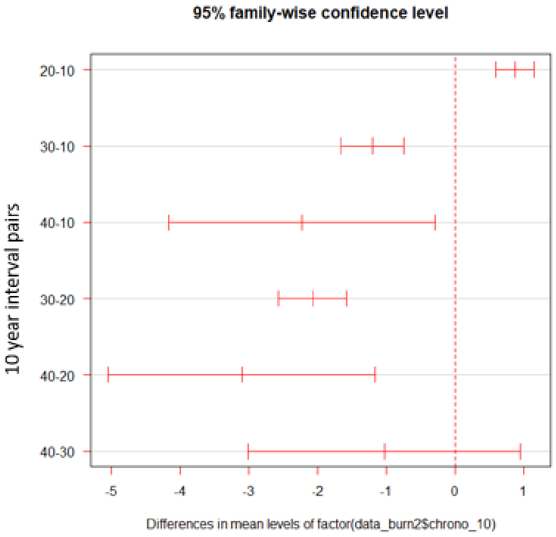

Northern Rockies

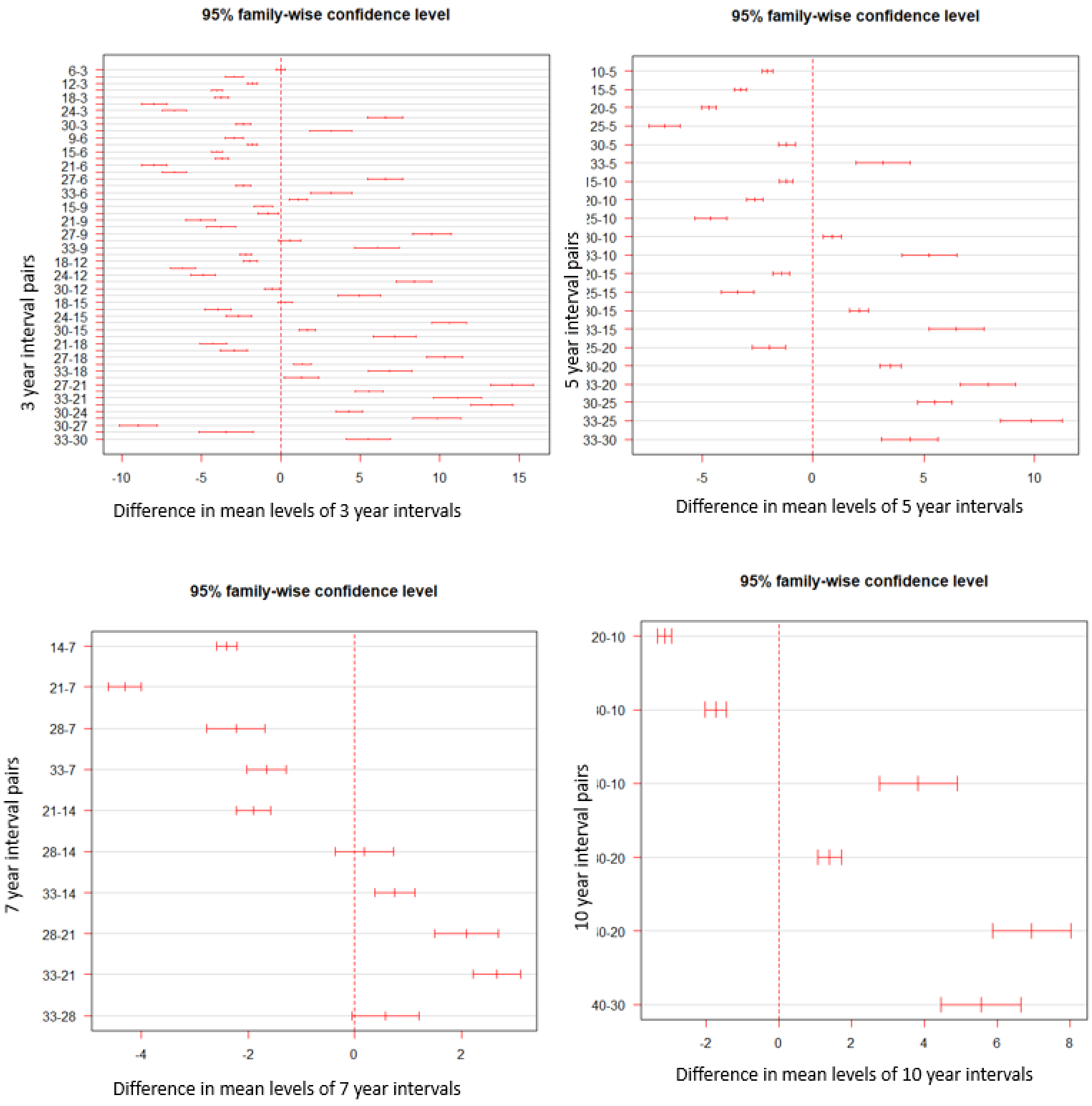

Southern Rockies

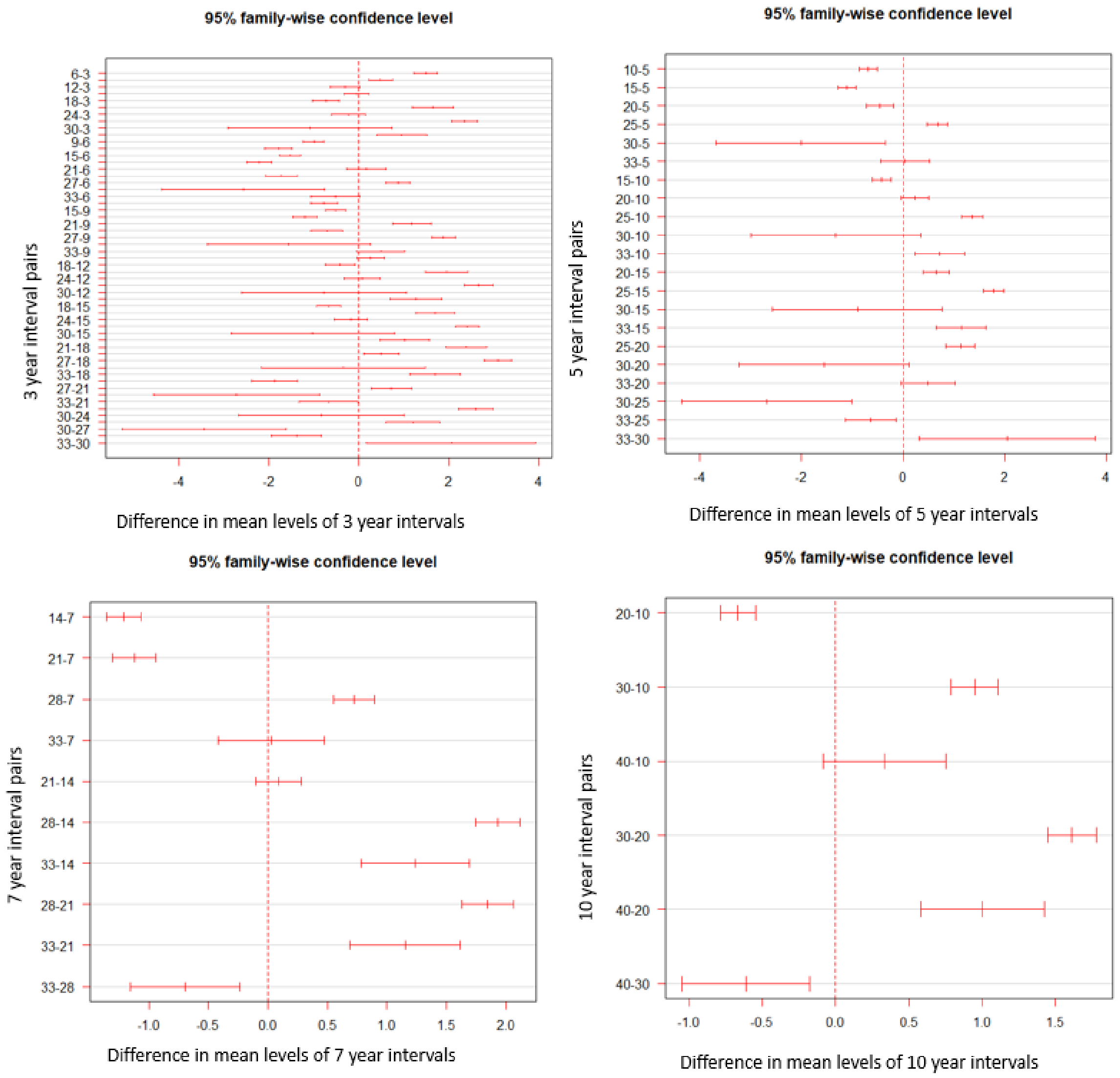


Supplementary Figure 4: Group mean and standard deviations of 3-, 5-, 7-, and 10- year intervals. A_ Pacific Northwest, b) Northern Rockies, and c) Southern Rockies.
